# The role of feature-based attention in visual serial dependence

**DOI:** 10.1101/584789

**Authors:** Matthias Fritsche, Floris P. de Lange

## Abstract

Perceptual decisions about current sensory input are biased towards input of the recent past – a phenomenon termed serial dependence. Serial dependence may serve to stabilize neural representations in the face of external and internal noise. However, it is unclear under which circumstances previous input attracts subsequent perceptual decisions, and thus, whether serial dependence reflects a broad smoothing or selective stabilization operation. Here, we investigated whether focusing attention on particular features of the previous stimulus modulates serial dependence. We found an attractive bias in orientation estimations when previous and current stimuli had similar orientations, and a repulsive bias when they had dissimilar orientations. The attractive bias was markedly reduced when observers attended to the size, rather than the orientation, of the previous stimulus. Conversely, the repulsive bias for stimuli with large orientation differences was not modulated by feature-based attention. This suggests separate sources of these positive and negative perceptual biases.

## Introduction

Humans often form perceptual decisions based on ambiguous and unstable sensory input. In vision, this instability is exacerbated by several factors such as eye movements, blinks and temporary occlusions of the visual scene. Moreover, further instabilities are introduced in the form of biological noise during neural processing. Yet, despite all these instabilities, we exhibit a remarkable capacity for making successful perceptual decisions. A key question is therefore how our brains maintain stable neural representations for perceptual decision making.

Importantly, our environment is relatively stable over short timescales and thus exhibits temporal continuity (Dong & Atick, 1995). Theoretically, this temporal continuity could be exploited to stabilize neural representations. In particular, by leveraging information from the recent past, neural representations could be smoothed in time to compensate for perturbations, which are not caused by genuine changes in the physical world (Burr & Cicchini, 2014). In line with this idea, recent studies have found that perceptual decisions about a large variety of visual stimulus features are biased towards features encountered in the recent past. Such features include orientation (Cicchini & Burr, 2017; Czoschke, Fischer, Beitner, Kaiser, & Bledowski, 2018; Fischer & Whitney, 2014; Fritsche et al., 2017), numerosity (Cicchini, Anobile, & Burr, 2014; Corbett, Fischer, & Whitney, 2011; Fornaciai & Park, 2018a), spatial location (Bliss, Sun, & D’Esposito, 2017; Manassi, Liberman, Kosovicheva, Zhang, & Whitney, 2018; Papadimitriou, White, & Snyder, 2016), visual variance (Suárez-Pinilla, Seth, & Roseboom, 2018), face identity (Liberman, Fischer, & Whitney, 2014), emotional expression (Liberman, Manassi, & Whitney, 2018) and attractiveness (Xia, Leib, & Whitney, 2016). While it is still debated whether such serial dependence biases are introduced at a perceptual stage or at a post-perceptual, decision or short-term memory stage (Fritsche et al, 2017; Bliss et al, 2017; Cicchini & Burr, 2017; Fornaciai & Park, 2018a), the ubiquity of serial dependencies in perceptual decisions is striking and suggests that serial dependencies might arise from a general computation of the brain, potentially reflecting the stabilization of neural representations.

Although serial dependence biases have been observed in perceptual decisions about a variety of stimulus features, the conditions under which they arise are still elusive. Consequently, the precise boundaries within which a stabilization of neural representations could take place are not known. In particular, it is unclear how a previous stimulus needs to be processed in order to exert a serial dependence bias in subsequent perceptual decisions. While serial dependence has been shown to depend on spatial attention towards the previous stimulus location (Fischer & Whitney, 2014), it has also been reported to occur when previous stimuli were task irrelevant (Fornaciai & Park, 2018a & b), suggesting that attention to a particular stimulus feature is not necessary for serial dependence to occur. In a similar vein, serial dependence is often thought to arise on the level of objects (Liberman et al., 2014, 2018), suggesting that attention to a particular stimulus feature might not be necessary as long as the object is spatially attended. Conversely, one recent study showed that serial dependence in judgments about visual variance only occurred when observers attended and reported the variance of a previous motion stimulus and not its direction (Suárez-Pinilla et al., 2018). While this result calls the independence of serial dependence from feature-based attention into question, it is unclear whether this would generalize to perceptual decisions about lower-level visual features such as orientation or numerosity. Furthermore, Suárez-Pinilla et al. employed different response methods for visual variance and direction reports, respectively, permitting the possibility that their results were influenced by serial dependencies in decisions about particular response adjustments, rather than about the stimulus feature itself. The current study, therefore, aims to elucidate the role of feature-based attention in serial dependence. To this end, we measured serial dependence in orientation estimations about gratings, while participants either attended to the orientation or size of a previous grating stimulus. Importantly, to isolate the effects of feature-based attention we tightly controlled the difficulty and the visual input in the two attention conditions.

Besides attractive serial dependence biases, previous studies have observed concurrent repulsive biases when subsequent stimuli differed markedly (Fritsche et al., 2017; Bliss et al., 2017). Currently, it is unclear whether attractive and repulsive biases originate from the same underlying neural process, or whether these are two distinct phenomena, concurrently observed in behavioral responses. To shed light on this question, we also assessed whether repulsive biases for stimuli with large orientation differences are modulated by feature-based attention. Different modulations of attractive and repulsive biases by feature-based attention would indicate that these biases arise from at least partially independent processes.

To preview, serial dependence biases in orientation judgments were strongly modulated by feature-based attention. That is, orientation estimations were biased towards the previous stimulus’ orientation, and this bias was twice as strong when participants attended the orientation versus the size of the previous stimulus. Strikingly, repulsive biases for stimuli with large orientation differences were also robustly present but not modulated by feature-based attention, suggesting that they may arise from an independent process, potentially akin to classical repulsive tilt-aftereffects (Gibson & Radner, 1937). Overall, the current study provides important insights into the conditions under which serial dependencies arise and demonstrates crucial boundaries within which stabilizations of neural representations through serial dependencies take place.

## Methods

### Participants

Thirty-eight naïve participants (27 female/11 male, age range 19 – 34 years) took part in the experiment. All participants reported normal or corrected-to-normal vision and gave written, informed consent prior to the start of the study. The study was approved by the local ethical review board (CMO region Arnhem-Nijmegen, The Netherlands) and was in accordance with the Declaration of Helsinki. Our target sample size was n = 34. This sample size was chosen to obtain 80% power for detecting a medium effect size (d = 0.5) with a two-sided paired t-test at an alpha level of 0.05. Four participants were excluded after the first experimental session due to insufficient performance and were not invited to the main experimental sessions. These participants were replaced with new participants to obtain 34 complete datasets. The experiment and analyses were preregistered on the Open Science Framework (https://osf.io/q7gj3/).

### Apparatus and stimuli

Visual stimuli were generated with the Psychophysics Toolbox (Brainard, 1997) for MATLAB (The MathWorks, Natick, MA) and were displayed on a 24′′ flat panel display (Benq XL2420T, resolution 1920 × 1080, refresh rate: 60 Hz). Participants viewed the stimuli from a distance of 53 cm in a dimly lit room, resting their head on a table-mounted chinrest.

A central white fixation dot of 0.25° visual angle diameter was presented on a mid-grey background throughout all experiment blocks. Participants were instructed to maintain fixation at all times. A cue stimulus, in the form of a white disc windowed by a Gaussian envelope (0.4° s.d.), was presented 9 visual degrees left or right from fixation. Reference stimuli were formed by a dark grey disc of variable size (3 – 3.5° radius) and two smaller, opposing discs (0.2° radius) of the same color that were offset 7° from the center of the reference stimulus. These smaller discs formed a reference orientation, which was defined as the orientation of a virtual line connecting the two small discs. Grating stimuli consisted of a sine wave grating (0.5 cycles/° spatial frequency, random phase, 8% Michelson contrast) with additive white noise smoothed with a 0.1° s.d. Gaussian kernel (16% contrast). Stimuli were masked with a circular aperture of variable size. The response bar stimulus was a white bar (0.4° width) windowed by a 1.2° s.d. Gaussian envelope and was presented at the same horizontal eccentricity as the cue and grating stimuli (**Fig. 1a** and **b**).

**Fig. 1.**
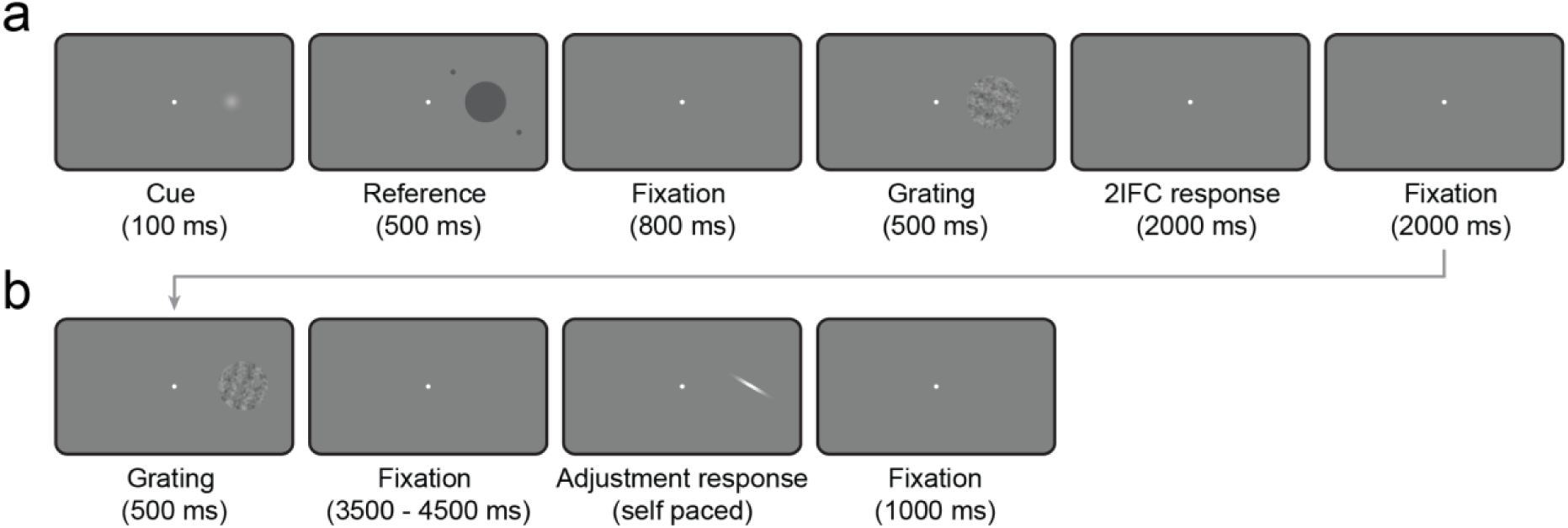
Sequence of events in the 2IFC (a) and main experiment tasks (a + b). In the 2IFC task (a) participants judged either the orientation or the size of a grating stimulus against a prior reference stimulus. The reference orientation was defined by a virtual line that connected the two outer discs of the reference stimulus, while the reference size was defined by the diameter of the inner disc. In separate blocks, participants either had to indicate whether the grating was oriented more clockwise or counterclockwise, or had a larger or smaller size than the reference. This task was employed in a staircase procedure to titrate the orientation and size differences between grating and reference to a set level of difficulty for the main experiment. In each trial of the main experiment (a + b), participants first performed either the orientation or size 2IFC judgment on the first grating stimulus (inducer), separated in different sessions. After the 2IFC response, participants saw a new grating stimulus (test) and subsequently reproduced the orientation of this test grating by adjusting a response bar. Stimulus presentation in the left or right visual field was pseudo-randomized across trials.

### Procedure

The experiment consisted of three separate sessions, each conducted on a different day, with consecutive sessions no more than two days apart. In the first session, we first measured each participant’s individual thresholds for the orientation and size two-interval forced choice (2IFC) tasks. In the remainder of the first session, participants practiced the serial dependence task, which they performed in the second and third session. Participants were not invited to the second and third sessions, if their orientation or size threshold for the 2IFC task exceeded priorly defined maximum thresholds, thereby excluding participants who were unable to perform the 2IFC tasks.

#### Threshold estimation

The sequence of events within each trial of the orientation and size 2IFC tasks is illustrated in **Figure 1a**. At the beginning of each trial a cue was presented left or right of fixation for 100 ms. After further 400 ms of fixation, a reference stimulus was presented on the same side as the cue for 500 ms. The inner disc of the reference stimulus had a radius randomly drawn from a uniform distribution on the interval [3, 3.5°]. The two outer discs of the reference stimulus formed a reference orientation, defined as the orientation of a virtual line connecting the two discs. On each trial, the reference orientation was randomly drawn from a uniform distribution on the interval of all possible orientations (0, 180°]. After 800 ms of fixation, a grating stimulus was presented on the same side as the cue and reference stimulus for 500 ms. The grating stimulus was oriented slightly more clockwise or counter-clockwise with respect to the reference orientation and was slightly smaller or larger than the inner disc of the reference stimulus. After the offset of the grating stimulus, there was a fixed 2000 ms response period and an inter-trial-interval of 2000 ms. Participants performed one of two tasks, separated into different blocks. In the orientation 2IFC task, participants judged whether the grating stimulus was oriented more clockwise or counter-clockwise than the reference orientation. In the size 2IFC task, participants judged whether the grating stimulus was smaller or larger than the reference disc. Responses were given via the arrow keys on a standard keyboard, and if no response was given within 2000 ms after the offset of the grating, the fixation dot briefly turned red and a new trial began. In order to avoid that participants focused on a subpart of the grating stimulus to solve the tasks, the spatial position of the grating stimulus was randomly jittered by a maximum of 1.5° visual angle on every trial.

The difficulty of the 2IFC tasks could be varied by changing the relative orientation *Δθ* (during the orientation task) and size *Δs* (during the size task) of the grating with respect to the reference stimulus. For each participant, we estimated their individual thresholds *Δθ* and *Δs* for performing at an accuracy of 75% on both tasks using the QUEST staircase algorithm (Watson & Pelli, 1983). We first estimated the size threshold *Δs*, while holding *Δθ* constant at ±10° orientation difference. Participants performed blocks of 48 trials. After each block the convergence of the *Δs* threshold estimate was visually inspected by the experimenter and the estimation was terminated after *Δs* converged to a stable value. Subsequently, we employed the same procedure to estimate the orientation threshold *Δθ*, while holding *Δs* constant at a ± 0.15° visual angle change in radius. Prior to each threshold estimation, participants performed one or more practice blocks with fixed values of *Δθ* = 15° and *Δs* = 0.3°, respectively, until they felt comfortable with the tasks. During the practice blocks, participants received feedback about the correctness of their response via brief color changes of the fixation dot to green (correct) or red (incorrect).

#### Serial dependence task

The sequence of events within each trial of the serial dependence task is illustrated in **Figure 1a** and **b**. The first part of each trial was identical to the 2IFC task, described above, with the exception that *Δθ* and *Δs* were now set to the individually estimated thresholds for each participant. Depending on session, participants either performed the orientation or the size 2IFC judgment. After the 2IFC response and 2000 ms of fixation, a second grating stimulus was presented for 500 ms at the same side as the cue, reference and first grating stimulus. This second grating had a relative orientation ranging from −90° to +90° in steps of 10° with respect to the first grating and all relative orientations occurred equally often in pseudorandomized order. After a variable delay ranging from 3500 to 4500 ms, a response bar with a random initial orientation appeared at the same location as the grating. Participants were asked to reproduce the orientation of the second grating by adjusting the response bar with the left and right arrow key. The response was submitted by pressing the space bar. The response was followed by a 1-second inter-trial-interval, before the next trial began. The role of the first grating stimulus, which was always compared in orientation or size to the reference stimulus, was to induce biases in adjustment responses to the subsequent second grating stimulus on each trial. Therefore, we term the first grating the “inducer” grating and the second grating the “test” grating. Since for the orientation 2IFC judgment participants had to focus on the orientation of the inducer and could neglect its size, we refer to the serial dependence task with the orientation 2IFC judgment as the “orientation” condition. Likewise, the serial dependence task with the size 2IFC judgment is termed “size” condition, as people had to focus on size and could ignore the inducer orientation.

Participants completed a total of 576 trials in the second and third session, each split into 8 blocks. Whether participants first performed the session with the orientation or size condition first was counterbalanced across participants. The horizontal location of the stimuli, the rotation and size change of the inducer grating with respect to the reference stimulus and the relative orientation of the test grating with respect to the inducer grating were pseudo-randomized across trials. Importantly, we used the exact same stimuli, trial parameters and trial sequence in both the second and third session, with the only difference that participants either judged the orientation or size difference of the inducer grating with respect to the reference.

### Data analysis

#### Outlier exclusion

We did not invite participants for the serial dependence task in the second and third session if their estimated orientation or size threshold in the first session exceeded *Δθ* = 20° or *Δs* = 0.4° visual angle, respectively. Furthermore, participants were excluded from data analysis if their thresholds for the orientation or size 2IFC task were more than 3 standard deviations above the group mean thresholds. Participants were also excluded if their mean 2IFC accuracy in either the orientation or size condition of the serial dependence task was below 60%. Finally, participants were excluded if their average response error in the orientation reproductions was more than 3 standard deviations above the group mean response error. According to these criteria, four participants were excluded after the first experimental session, because their *Δθ* threshold in the orientation 2IFC task exceeded 20°.

For the analysis of serial dependence biases, we excluded individual trials for which the absolute adjustment error was more than 3 standard deviations away from the average adjustment error of that participant. Furthermore, we excluded trials in which no 2IFC response was given within the 2000 ms response period. Prior to further analyses, we removed each participant’s mean adjustment response error from the adjustment data, separately for each attention condition to remove general clockwise or counterclockwise response biases that are independent of biases due to stimulus history. On average, we rejected 5.62 of 576 trials per participant due to outlier adjustment responses (orientation condition: M = 2.79, SE = 0.35; size condition: M = 2.82, SE = 0.31). Furthermore, we rejected an average of 8.26 trials per participant because no 2IFC response was given within the 2000 ms response period (orientation condition: M = 3.88, SE = 0.90; size condition: M = 4.38, SE = 0.66).

#### Accuracies and response times

We statistically compared the 2IFC accuracies and the mean adjustment errors in the orientation and size conditions of the serial dependence task with a two-sided paired t-test and a Bayesian undirected paired-sample t-test with a Cauchy prior with a default scale of 0.707 (Rouder, Speckman, Sun, Morey & Iverson, 2009). Similarly, we assessed potential differences in response times in both the 2IFC and adjustment responses across the two conditions.

#### Serial dependence on inducer stimulus

In order to quantify systematic biases in adjustment responses about the test grating towards the orientation of the inducer grating, i.e. serial dependence, we first expressed the adjustment response errors as a function of the orientation difference between inducer and test gratings. For positive values of this orientation difference, the inducer grating was oriented more clockwise than the test grating. Similarly, positive response errors denote trials in which the response bar was adjusted more clockwise than the test grating. Next, we pooled the response errors of all participants, in the orientation and size conditions respectively, and fitted derivative of a Gaussian curves (DoG) to the group data in both conditions. The DoG is given by 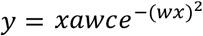, where x is the relative orientation of the inducer grating, *a* is the amplitude of the curve peaks, *w* is the width of the curve and *c* is the constant 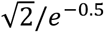. The constant *c* is chosen such that parameter *a* numerically matches the height of the curve peak. The amplitude parameter *a* was taken as the strength of serial dependence, as it indicates how much the response to the test orientation could be biased towards the inducer orientation for the maximally effective orientation difference between stimuli. For all model fits, the width parameter *w* of the DoG curve was treated as a free parameter, constrained to a range of plausible values (w = 0.02 – 0.07, corresponding to curve peaks between 10° and 35° orientation difference).

We used permutation tests to statistically assess serial dependence biases in the orientation and size conditions. A single permutation was computed by first randomly inverting the signs of each participant’s response errors (i.e. changing the direction of the response errors). This is equivalent to randomly shuffling the labels between the empirically observed data and a distribution of no serial dependence (a flat surrogate response error distribution) and subtracting the two conditions from each other per participant. Subsequently, we fit a new DoG model to the pooled group data and collected the resulting amplitude parameter in a permutation distribution. We repeated this permutation procedure 10,000 times. As p-values we report the percentage of permutations that led to equal or more extreme values for the amplitude parameter than the one estimated on the empirical data. The significance level was set to α = 0.05 (one-sided permutation test) for testing serial dependence in the orientation condition, as we expected to find attraction biases, in line with previous studies. The significance level for the size condition was set to α = 0.025 (two-sided permutation test), since due to the lack of previous experimental evidence we regarded both attraction and repulsion biases as possible. The exchangeability requirement for permutation tests is met, because under the null hypothesis of no serial dependence, the labels of the empirically observed data and a flat surrogate response error distribution of no serial dependence are exchangeable.

In order to assess the difference in serial dependence between the orientation and size condition we employed a permutation test as well. For each permutation, we randomly shuffled the condition labels of the orientation and size condition of each participant. We then fit DoG models to the permuted conditions, pooled across participants, and recorded the difference between the amplitude parameters. We repeated this procedure 10,000 times. As p-values we reported the percentage of permutations that led to an equal or more extreme amplitude difference than the one we observed in the experiment. The significance level was set to α = 0.025 (two-sided permutation test). The exchangeability requirement for permutation tests is met, because under the null hypothesis of no difference in serial dependence between the orientation and size conditions, the condition labels are exchangeable.

In addition to this analysis in which we pooled data across participants, we conducted a second analysis in which we fit DoG models to the data of each participant, thereby obtaining estimates of serial dependence biases for each individual. We compared the group’s mean bias in each condition to zero using one-sample t-tests. As with the permutation tests, we used a one-sided t-test in the orientation condition and a two-sided t-test in the size condition. To assess a difference in biases across conditions we employed a two-sided paired t-test. All significance levels were set to α = 0.05. Furthermore, for all classical null-hypothesis significance tests we conducted the analogous Bayesian t-tests with default Cauchy priors (scale 0.707).

#### Repulsive biases for large orientation differences

Next to positive attraction biases for subsequent stimuli with similar orientations, we expected to observe negative repulsive biases for stimuli with large orientation differences beyond 60° (Bliss et al.; Fritsche et al., 2017). In order to test whether such repulsive biases also occurred in the current experiment, we averaged each participant’s adjustment response errors in a negative and positive bin. The negative bin comprised trials with −80, −70 and −60° orientation difference between inducer and test grating, whereas the positive bin comprised trials with 60, 70 and 80° orientation difference. For each participant, we computed a bias by subtracting the average response error in the negative bin from the error in the positive bin and dividing the resulting value by two. Negative values for this bias reflect repulsive biases of adjustment responses away from the inducer stimulus. We statistically compared the biases in the orientation and size condition against zero using one-sample t-tests. Analogous to the tests of positive serial dependence, we employed a one-sided test in the orientation condition and a two-sided test in the size condition at significance levels of α = 0.05. Moreover, we were interested whether the repulsive biases for large orientation differences were modulated by feature-based attention towards the orientation or size of the inducer stimulus. To this end, we compared the biases in the orientation and size conditions using a two-sided paired t-test. Similar, to the analyses above we also conducted Bayesian t-tests with default Cauchy priors (scale 0.707).

#### Serial dependence on previous test stimulus

The current experiment was primarily designed to measure orientation estimation biases towards or away from a preceding inducer stimulus, which was either attended in terms of its orientation or size. However, previous studies not only found attractive biases towards immediately preceding stimuli, but also to stimuli seen further in the past (Fischer & Whitney, 2014; Fritsche et al., 2017). Two interesting questions derive from these previous findings in the context of the current study. First, we wondered whether we could replicate the finding of attractive serial dependence biases towards temporally more distant stimuli. Second, we asked whether feature-based attention on the inducer stimulus would not only modulate the attraction bias towards this inducer, but also modulate the attraction bias to stimuli that were presented further in the past. In other words, is the serial dependence bias towards the recent stimulus history modulated by how intervening information is processed? To shed light on this question we conducted a further exploratory analysis that was not part of the preregistered analysis plan. In this analysis we investigated whether adjustment responses were not only biased towards the inducer stimulus on the same trial, but also towards the test stimulus presented on the previous trial (Fritsche et al., 2017). To this end, we repeated the serial dependence analysis described above, with the exception that response errors were now expressed as a function of the orientation difference between the test stimuli presented on the previous and current trial, respectively. Since we expected the attraction biases towards previous trial’s test stimulus to be weaker and more variable, potentially resulting in problems with model fits to single subject data, we focused on fitting the DoG models to the group data and statistically assessed the amplitude estimates with the random-effects permutation test described above. Notably, the orientation and size condition of the experiment were similar in the sense that participants always attended to the orientation of each test stimulus, as they had to reproduce its orientation. However, the conditions differed in the processing of the inducer stimulus which was presented in between the previous and current test stimulus and was either attended in terms of orientation or size. Importantly, if information from different stimuli encountered in the recent past would be integrated independently with the current stimulus representation, then this differential processing of the intervening inducer stimulus should not impact biases between test stimuli of successive trials. Conversely, if stimulus information from different moments in the recent history would interact or interfere, attending to the orientation or size of an intervening inducer stimulus could influence serial dependencies between successive test stimuli.

### Results

#### Overall task performance

The accuracies of the 2IFC judgments in the main task were close to the target accuracy of 75% (orientation condition: M = 74.70%, SE = 0.81%; size condition: M = 77.03%, SE = 0.81%), and the average thresholds were *Δθ* = 10.17° (SE = 0.64°) and *Δs* = 0.13° visual angle (SE = 0.006°), respectively. While a paired t-test revealed that participants performed significantly more accurate in the size 2IFC task (t(33) = −2.33, p = 0.03), a Bayesian t-test indicated only anecdotal evidence for a difference in 2IFC performance across conditions (BF_10_ = 1.93). Similarly, the average error in adjustment responses was slightly but significantly lower in the size compared to the orientation condition (orientation condition: M = 8.99°, SE = 0.30°; size condition: M = 8.61°, SE = 0.31°; difference: t(33) = 2.41, p = 0.02), while a Bayesian t-test again only indicated anecdotal evidence for a difference across conditions (BF_10_ = 2.27). We observed that participants gave significantly faster 2IFC responses in the size than in the orientation condition (orientation condition: M = 0.60 seconds, SE = 0.03 seconds; size condition: M = 0.49 seconds, SE = 0.03 seconds; difference: t(33) = 4.40, p < 0.001, BF_10_ = 240). We note that despite this difference in responses times, the response period was always 2000 ms, and therefore there was no difference in inter-stimulus intervals between inducer and test stimuli across conditions. Finally, there was no significant difference between adjustment response times across conditions (orientation condition: M = 2.45 seconds, SE = 0.10 seconds; size condition: M = 2.45, SE = 0.08 seconds; difference: t(33) = 0.04, p = 0.97, BF_01_ = 5.44).

#### Serial dependence on inducer stimulus is modulated by feature-based attention

Adjustment responses to test stimuli were systematically attracted towards the orientation of the preceding inducer stimulus when it was of similar orientation (**Fig. 2a** and **b**), both in the orientation condition (*a* = 2.36°, p < 0.0001, permutation test) and size condition (*a* = 1.01°, p < 0.0001, permutation test). Crucially however, this serial dependence bias was significantly stronger when people attended to the orientation of the inducer stimulus, compared to when they attended to its size (p < 0.0001, permutation test). Furthermore, the peak locations of the DoG model were significantly more narrow in the size compared to the orientation condition (±13.74° vs. ±17.88° peak locations, p = 0.01, permutation test, not preregistered). These results were corroborated by a second analysis in which we fitted DoG models to the individual participant data. This complementary analysis revealed significant serial dependence both when participants were attending to orientation (amplitude *a*: M = 2.60°, SE = 0.38, t(33) = 6.73, p < 0.0001, BF_+1_ = 2.6e+5) and size of the inducer (amplitude *a:* M = 0.81°, SE = 0.29°, t(33) = 2.77, p = 0.009, BF_10_ = 4.64). Similar to the first analysis serial dependence was significantly stronger when attending the orientation of the inducer (t(33) = 4.74, p < 0.0001, BF_10_ = 578). To summarize, while adjustment responses were biased towards the inducer orientation, even when the inducer orientation was not attended, the attraction bias was more than twice as strong when the inducer orientation was attended. Therefore, positive serial dependence is strongly modulated by feature-based attention towards previous stimulus features.

**Fig. 2.**
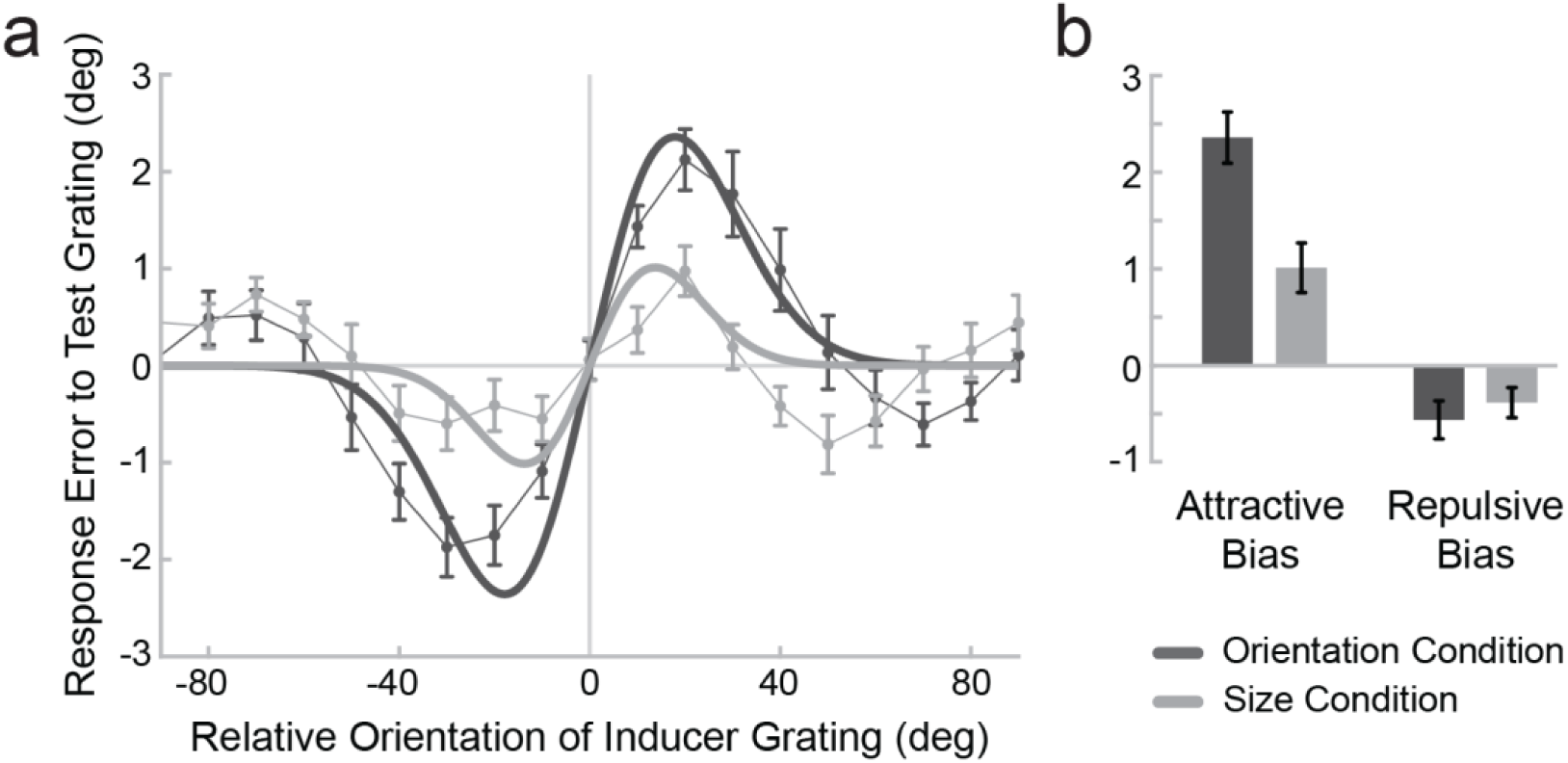
Systematic attractive and repulsive biases of adjustment responses to the inducer grating. (a) Serial dependence error plot of trials in which participants attended to the orientation (dark gray) or size (light gray) of the inducer. We expressed the adjustment response errors (y axis) as a function of the orientation difference between inducer and test grating (x axis). For positive x values the inducer was oriented more clockwise than the test grating and for positive y values the current response error was in the clockwise direction. Responses to test gratings with orientations similar to the inducer were systematically biased towards the inducer, as can be seen from the group average response errors (dark and light gray data points; data points were smoothed by averaging with the respective neighboring data points). This bias follows a Derivative-of-Gaussian shape (DoG, model fits shown as think dark and light gray lines).

Crucially, the magnitude of this attractive bias was significantly weaker, when participants judged the size, instead of the orientation of the inducer stimulus (b, left), as indicated by a reduced amplitude parameter of the DoG model (error bars show bootstrapped SEMs). Additionally, next to the attractive bias between inducer and test grating with similar orientations, adjustment responses were repelled from the inducer when inducer and test grating had very different orientations (difference ≥ 60°). Notably, unlike the attractive bias this repulsive bias appeared not to be modulated by attending to either the orientation or size of the inducer (b, right). Error bars depict SEMs.

#### Repulsive biases for large orientation differences are not modulated by feature-based attention

Next to the attraction bias towards previous inducer stimuli with similar orientations, adjustment responses were repelled away from inducer stimuli with large orientation differences (**Fig. 2b**). This held true both when participants attended the orientation (t(33) = −2.81, p = 0.004, BF-_1_ = 10) and the size of the inducer stimulus (t(33) = −2.43, p = 0.02, BF_10_ = 2.33). However, there was no significant difference in repulsion biases across the two conditions (t(33) = −0.74, p = 0.46) and a Bayesian t-test revealed moderate evidence for the null hypothesis of no difference across conditions (BF_01_ = 4.22). This finding indicates that, unlike the positive serial dependencies between stimuli with similar orientations, the repulsive biases for successive stimuli with large orientation differences are not modulated by feature-based attention. This suggests that the attractive and repulsive biases measured in adjustment responses may originate, at least partly, from separate underlying processes and are superimposed in the final behavioral response.

One may wonder whether the current finding that repulsive biases for successive stimuli with large orientation differences are not modulated by feature-based attention might critically depend on the pre-registered analysis choice of bin widths from ±60 to ±80°, over which the repulsive biases were computed. This concern is strengthened by the observation that feature-based attention appears to not only modulate the amplitude of the positive serial dependence bias, but also its width (**Fig. 2a**). As a consequence, manipulating positive biases in the center of the serial dependence plot may have a systematic impact on the repulsive biases expressed in the periphery. For instance, a narrower tuning of the central positive biases might be accompanied by a shift of the repulsive biases towards the center of the error plot. In this case, computing the repulsive biases within fixed peripheral bins might underestimate the actual magnitude of the repulsion. Consequently, comparing the repulsive biases over the same orientation differences in both attention conditions might lead to a biased estimation of the repulsive biases. In order to overcome this potential issue, we conducted an additional exploratory control analysis of the repulsive biases, following a multiverse analysis approach (Steegen, Tuerlinckx, Gelman & Vanpaemel, 2016). This control analysis was similar to our original analysis, with the exception that we allowed the bin widths for which we computed the repulsive biases to vary independently in the orientation and size conditions. Specifically, we varied the number of bins included in the analysis by in-or excluding bins towards the center of the serial dependence plot, i.e. computing biases over the range [±X, ±80], where X was varied between 40 and 80° in steps of 10°. As a result, we could compare biases in the orientation condition that were, for instance, computed over a range from ±60 to ±80° orientation difference to biases in the size condition that were computed from ±40 to ±80° orientation difference, thereby investigating the robustness of the current result in light of different analysis choices for bin widths (see **Fig. S1**). The analysis revealed significant repulsion biases for X ≥ 60° in the orientation condition, and for 40° ≤ X ≤ 60° in the size condition (all p < 0.05). Moreover, for those values of X for which there were significant repulsion biases in both conditions, there was no significant difference between biases across conditions (all p > 0.25). Similarly, Bayesian t-test revealed moderate evidence for the null hypothesis of no difference across conditions for all but one comparison (all BF_01_ > 3, except X_ori_ = 80°, X_size_ = 40° for which BF_01_ = 2.91). To conclude, even when computing repulsive biases over a wide range of variable orientation differences, there is no evidence that feature-based attention modulates these repulsive biases.

#### Serial dependence on previous test stimulus is modulated by attention to intervening inducer

In an additional exploratory analysis, we investigated whether adjustment responses to the current test stimulus were also systematically attracted towards the previous test stimulus, as has been reported before (Cicchini et al., 2017; Fritsche et al., 2017). Of particular interest, we explored whether this attraction bias was modulated by *how* the intervening inducer stimulus was processed, in terms of feature-based attention to the intervening inducer stimulus. Although there was always an inducer stimulus presented between test stimuli of subsequent trials, previous studies reported that serial dependencies can exist for stimuli seen up to 10 - 15 seconds in the past (Fischer & Whitney, 2014). In the current experiment, the onset of a new test stimulus occurred on average ~13.4 seconds after the offset of the previous test stimulus, and ~6.9 seconds after the offset of the previous adjustment response. Indeed, we found that adjustment responses were significantly biased towards the test stimulus on the previous trial (**Fig. 3**), both in the orientation condition (*a* = 1.38°, p < 0.0001, permutation test) and the size condition (*a* = 2.18°, p < 0.0001, permutation test). Surprisingly, the bias was significantly stronger in the size than in the orientation condition (p = 0.0003, permutation test). That is, although the previous test stimulus was processed similarly in both conditions, serial dependence was stronger when participants attended to the size of an intervening inducer stimulus, compared to when the focused on its orientation. To our knowledge, this is the first indication that positive serial dependencies for previous stimuli are modulated by how intervening stimuli are processed. This might provide an important constraint for computational models of serial dependence biases, as it suggests that representations of the recent perceptual history interact or interfere and are not independently integrated with the current stimulus representation.

**Fig. 3.**
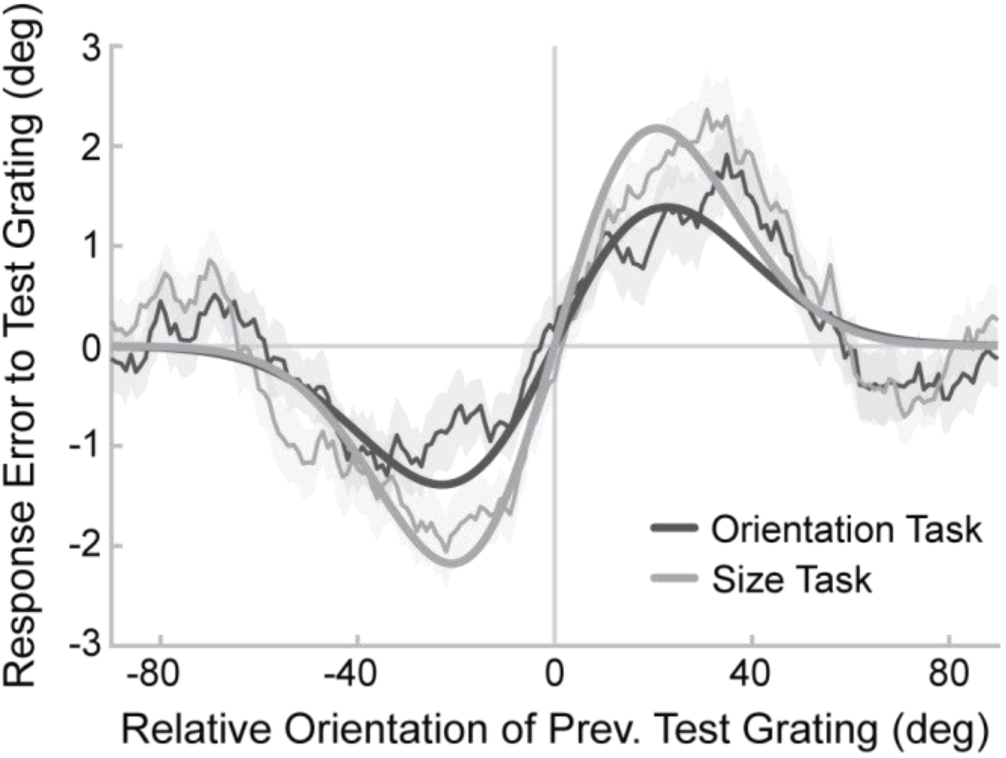
Serial dependence bias towards test grating on previous trial is modulated by attention to intervening inducer. We assessed serial dependence towards the previous test grating similar as in Fig. 2a, but conditioned on the relative orientation of the previous test grating, instead of the current trial’s inducer. Intriguingly, the attraction bias towards the test grating of the previous trial was stronger when participants judged the size, instead of the orientation, of the intervening inducer grating (p = 0.0003).

## Discussion

We have shown that attractive serial dependence biases in orientation estimations are strongly modulated by feature-based attention. That is, orientation estimations of a stimulus are more strongly attracted to the orientation of a previous inducer grating when observers attended to the inducer’s orientation rather than its size. Importantly, in both cases observers attended the same object at the same spatial location, precluding a difference in serial dependence due to differences in object-based or spatial attention (Fischer & Whitney, 2014). Integrating the current and previous findings on the role of attention in positive serial dependence suggests a very selective smoothing operation, which limits the role of serial dependence in stabilizing perceptual representations within the entire visual scene. As such, it is unlikely that serial dependence plays a major role in phenomena such as change blindness (Burr & Cicchini, 2014; Rensink, O’Regan, & Clark, 1997; Simons & Levin, 1997), which are marked by the absence of attention to a missed feature in a visual scene. Furthermore, the current finding that serial dependence can selectively smooth attended features of an object over time while leaving unattended features of the same object relatively unsmoothed suggests that it may, in part, operate below the level of object representations.

Interestingly, attractive serial dependence biases in orientation estimations did not completely disappear when observers attended to the size of the inducer stimulus. It is possible that the remaining small but robust attraction bias reflects a distinct bias, which is attention-independent, potentially arising from different processes than the attention-dependent bias. However, a perhaps more plausible explanation is that even though observers were asked to focus only on the size in the size 2IFC task, they might have nevertheless paid some attention to orientation as well, leading to small attraction biases in the size condition. In a similar vein, previous findings of serial dependence to task irrelevant stimulus features could be explained by residual attention to the feature dimension for which serial dependence was assessed (Fornaciai & Park, 2018a & b).

It is worthwhile to note that in the current experiment attractive serial dependence in orientation reproductions occurred even though the inducer orientation was never reproduced, but only judged with respect to a reference stimulus. Importantly, this judgment of the inducer’s orientation with respect to the reference was independent with respect to the orientation of the subsequent test stimulus. This provides corroborating evidence that an adjustment response, or the covert preparation thereof, is not necessary for inducing serial dependence biases, confirming previous claims (Fischer & Whitney, 2014).

Apart from the attraction biases for inducer and test stimuli with similar orientations, we found that adjustment responses were repelled when inducer and test stimuli had markedly different orientations (Bliss et al., 2018; Fritsche et al., 2017). Strikingly, we found no evidence for a modulation of these repulsive biases by feature-based attention. This asymmetric influence of attention on attractive and repulsive biases suggests that the two biases might originate from, at least partially, independent neural processes, and cautions against devising computational models of serial dependence in which positive and negative biases are co-dependent. These observations are also in line with a previous finding that the repulsive biases for successive stimuli with large orientation differences disappeared when presenting stimuli at different spatial locations, whereas the attractive bias remained of identical magnitude (Fritsche et al., 2017). Together, these findings suggest that the repulsive biases might occur at an early, retinotopic stage, which is not strongly modulated by attention, while the attraction bias might occur at a later, more spatially invariant stage, which is strongly affected by feature-based attention. However, the exact nature of the repulsive biases remains elusive. While reminiscent of classical negative perceptual adaptation (Webster, 2015), classical tilt-aftereffects predominantly occur for orientation differences between 0 and 45° and can turn into attractive biases for larger differences (Gibson & Radner, 1937). Furthermore, perceptual adaptation effects have been found to be modulated, albeit weakly, by feature-based attention (Spivey & Spirn, 2000; Kreutzer, Fink, & Weidner, 2015). Thus, it appears unlikely that the currently observed repulsive biases reflect classical perceptual adaptation. However, perceptual adaptation effects can occur even when inducer stimuli are rendered invisible by crowding (He, Cavanagh & Intrilligator, 1996), binocular rivalry (Wade & Wenderoth, 1978) and continuous flash suppression (Maruya, Watanabe & Watanabe, 2008) and neural adaptation can be observed in anesthetized animals (Kohn & Movshon, 2003), suggesting that adaptation can occur in the absence of attentional selection. Therefore, whether the present repulsive biases reflect a form of classical adaptation or a distinct phenomenon will be an interesting topic for future research.

Finally, in an exploratory analysis we asked whether the attractive serial dependence bias towards the recent stimulus history is modulated by how intervening information is processed. While it has been shown previously that attractive serial dependence exists beyond just the immediately preceding stimulus (Fischer & Whitney, 2014), the way in which multiple previous stimuli jointly lead to a serial dependence bias in the current estimation response is not known and computational models accounting for this long-term serial dependencies are scarce (but see Kalm & Norris, 2018). Surprisingly, we found that orientation estimations were more strongly biased towards test stimuli of the previous trial, when observers attended to size and not orientation of the intervening inducer stimuli. Crucially, one would not expect such an influence of the processing of intervening stimuli, if representations of recently encountered stimuli were independently integrated with the current stimulus representation. In turn, to a first approximation this suggests that representations of the recent perceptual history interact or interfere when maintained in memory and integrated with new information. Unfortunately, the current experimental design, with interleaved adjustment and 2IFC responses, is not ideally suited to quantitatively assess computational models of serial dependence, such as a recently proposed mixture model of internal representations (Kalm & Norris, 2018). However, the current exploratory observation may present an important feature of serial dependence that would need to be accounted for in computational models of this phenomenon.

To conclude, we have demonstrated that attractive serial dependence in orientation estimations is strongly modulated by feature-based attention, while repulsive biases for large orientation differences are not. This presents a distinguishing feature for positive and negative biases that are concurrently observed in perceptual estimations. Furthermore, our findings provide important insights into the conditions under which attractive serial dependencies occur, indicating a selective smoothing operation, which stabilizes representations of successively attended features of the same kind. The current study therefore contributes to understanding the boundaries within which stabilization of neural representations through serial dependence can take place.

## Acknowledgements

We thank Joey Zhou and Eelke Spaak for comments on the manuscript. This work was supported by a grant from the European Union Horizon 2020 Program (ERC Starting Grant 678286, “Contextvision”).

## Author contributions

Both authors designed the study. M.F. recorded and analyzed the data. Both authors wrote the manuscript.

## Supplemental Material

**Fig. S1.**
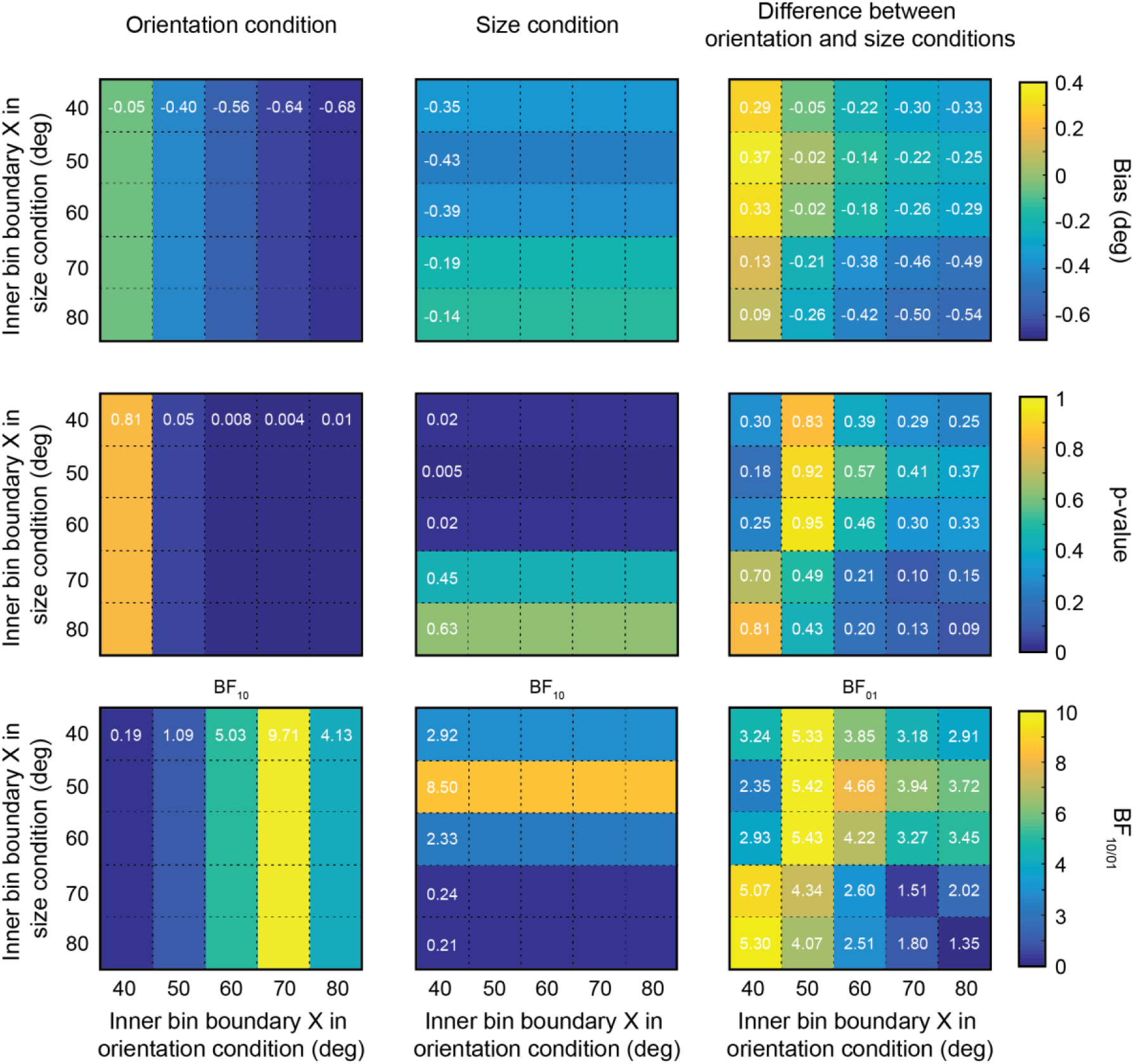
Exploratory control analysis of repulsive biases for large orientation differences between inducer and test stimuli. Biases were computed by averaging response errors in orientation bins ranging from ±X to ±80°, where X was varied from ±40 to ±80°. Biases from negative and positive orientation bins were combined by subtracting the bias in the negative bin from the positive bin and dividing the resulting bias by two. Negative values indicate repulsive biases. The upper row shows the bias magnitudes in the orientation (left) and size conditions (middle) and their difference (right). The middle row shows p-values of one-sample t-tests against zero (left/middle), and a paired t-test between the conditions (right). The lower row shows analogous Bayesian t-tests. Left and middle panels show BF_10_, i.e. the evidence for the alternative hypothesis of a bias different from zero versus the null hypothesis of no bias. The right panel shows BF_01_, i.e. the evidence for the null hypothesis of no difference between conditions. One can see that there are significant repulsion biases for X ≥ 60° in the orientation condition, and for 40° ≤ X ≤ 60° in the size condition. Furthermore, for these values of X for which there were significant repulsion biases in both conditions, there was no significant difference between biases across conditions (all p > 0.25, not corrected for multiple comparisons). Similarly, Bayesian t-test revealed moderate evidence for the null hypothesis of no difference across conditions for all but one comparison (all BF_01_ > 3, except X_ori_ = 80°, X_size_ = 40° for which BF_01_ = 2.911). Therefore, our finding that repulsive biases for large orientation differences are not modulated by feature-based attention was not specific to the particular bin width specifications that were preregistered, but extends to widely variable bin sizes.

